# Conservation and evolution of the programmed ribosomal frameshift in *prfB* across the bacterial domain

**DOI:** 10.1101/2024.09.24.614795

**Authors:** Cassidy R. Prince, Isabella N. Lin, Heather A. Feaga

## Abstract

When the ribosome reaches a stop codon, translation is terminated by a release factor. Bacteria encode two release factors, RF1 and RF2. In many bacteria, the gene encoding RF2 (*prfB)* contains an in-frame premature stop codon near the beginning of the open reading frame. A programmed ribosomal frameshift is therefore required to translate full-length RF2. While the molecular mechanism of the programmed ribosomal frameshift has been extensively characterized in *Escherichia coli*, bioinformatic analysis of the evolution and conservation of this motif has been limited to few genomes. By analyzing >12,000 bacterial genomes, we sought to thoroughly characterize the conserved frameshifting elements within the programmed frameshifting motif and identify genomic features of phyla that have lost the motif altogether. We find that the programmed ribosomal frameshift in *prfB* was likely present in the last common ancestor of bacteria and that the motif elements are almost completely conserved, including the identity of the internal stop codon. We find that loss of the programmed frameshift motif is highly correlated with RF2-specific stop codon usage, suggesting that stop codon usage has shaped the conservation of this regulatory mechanism. In support of this model, the programmed frameshift in *prfB* is entirely absent in Actinobacteriota, which have particularly high RF2 specific stop codon usage. Finally, we show that a model member of Actinobacteriota fails to produce full-length RF2 when provided with an allele of *prfB* that contains the programmed frameshifting motif. Altogether, our work provides a thorough characterization of RF2 regulation across the bacterial domain.

## Introduction

Ribosomes translate mRNA into protein by iteratively decoding one codon at a time. Reading frame maintenance is essential for translation fidelity. Frameshifting alters the identity of downstream amino acids and typically leads to premature termination since the ribosome quickly encounters a stop codon when it deviates from an open reading frame (1). In rare cases, ribosomal frameshifting is required to make a full-length protein (2–5). These types of frameshifts are referred to as programmed ribosomal frameshifts (6–9)

A textbook example of programmed ribosomal frameshifting in bacteria is found in *prfB*, the gene that encodes Release Factor 2 (RF2) (10, 11). RF2 terminates translation at UGA and UAA stop codons (12). In *Escherichia coli prfB*, there is an in-frame UGA codon at position 26 of the open reading frame (10) (Fig. 1). A purine rich, internal Shine-Dalgarno (SD)-like sequence is located 6 nucleotides upstream of this stop codon and a CTT leucine codon immediately precedes the stop codon (13). The UGA stop codon is followed by a C nucleotide, making it a weak stop codon (14–16). *In vitro* and *in vivo* studies using *E. coli* show that when concentrations of RF2 are high, RF2 terminates translation at the premature stop codon, creating a truncated peptide (11, 17). When RF2 concentrations are low, the ribosome pauses at the stop codon, and the SD-like sequence in the mRNA base-pairs with the anti-SD at the 3’ end of the 16S rRNA, displacing the E-site tRNA (13, 17, 18). The leucine tRNA in the P site then slips into the +1 frame, causing the ribosome to bypass the internal stop codon and continue translation to produce full-length RF2 (11, 13). Since RF2 levels directly regulate the translation of the *prfB* transcript, this frameshifting mechanism autoregulates RF2 levels.

**Figure 1.**
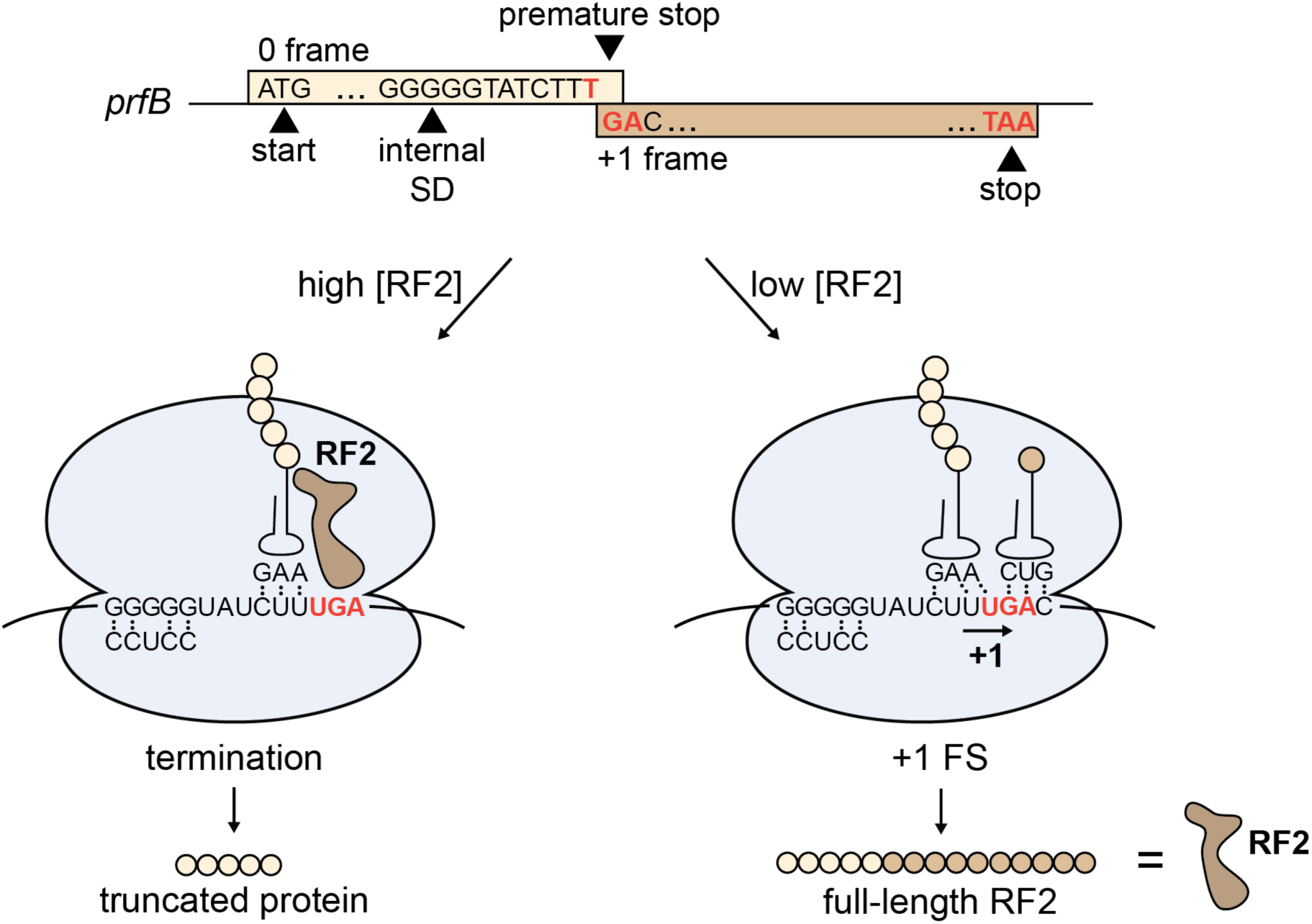
Schematic of *prfB* programmed ribosomal frameshift mechanism. (Top) DNA sequence elements of the *prfB* gene sequence in *E. coli*. (Bottom) The ribosome pauses at the internal SD-like sequence GGGGG, which base-pairs with the anti-SD sequence on the small ribosomal subunit. If RF2 levels are high, RF2 will enter the A-site and terminate translation at the premature UGA stop codon. If RF2 levels are low, the ribosome will slip into the +1 reading frame, causing a GAC codon to be positioned in the A-site instead of the premature stop codon. Therefore, ribosomal frameshifting results in production of full-length RF2.

The first bioinformatic analysis of the conservation of the frameshifting motif was performed by Baranov and colleagues in 2002, with 87 bacterial *prfB* sequences (19). This foundational work determined that 70% of these *prfB* sequences required the programmed ribosomal frameshift to produce RF2, and that in all but one sequence, the in-frame stop codon interrupting the *prfB* open reading frame was UGA. The slippery sequence (CTT) and the C nucleotide immediately following the stop codon were all highly conserved (20). Follow-up on a larger set of 259 genomes uncovered the regulatory frameshift in 87% of these genomes, indicating that the prevalence of this frameshift was even more widespread than previously determined (21).

Tens of thousands of bacterial genomes are now available, facilitating more thorough analyses of the conservation of the *prfB* programmed frameshift as an auto-regulatory mechanism. Our analyses are also aided by recent developments in bacterial phylogeny, which better resolve the bacterial species tree. Using a diverse and comprehensive genome set, we sought to answer remaining questions about the evolution and conservation of the programmed frameshift autoregulatory mechanism in *prfB*. For example, was the programmed frameshift present in the ancestral *prfB*, and what are the genome characteristics that are associated with absence of the programmed frameshift?

We surveyed 12,751 bacterial genomes across 21 phyla to determine the prevalence and conserved features of programmed ribosomal frameshifting in *prfB*. We find that most bacterial genomes encode a *prfB* that requires a ribosomal frameshift to produce full-length RF2 and that the programmed frameshift was most likely present in the common ancestor of bacteria. In our dataset, the frameshift sequence motif is extremely well-conserved, and most strikingly, the identity of the premature stop codon is nearly always an RF2-specific UGA stop codon. We did not find any instance of the RF1-specific stop codon within the motif, indicating that the autoregulatory role of the programmed ribosomal frameshift in *prfB* is conserved across the bacterial domain. Next, we examined the species that lack the autoregulatory frameshift motif and found that they have significantly higher RF2 stop codon usage, which may explain why RF2 is not autoregulated by this mechanism in these organisms. Consistent with this model, the Actinobacterium *Mycobacterium smegmatis*, which has high RF2 stop codon usage, exhibits low frameshifting efficiency at the motif. Cumulatively, our results support the autoregulatory function of the *prfB* frameshift across the bacterial domain, suggest that this autoregulatory mechanism was present in ancestral *prfB*, and that this autoregulation was lost in phyla that exhibit high RF2-specific stop codon usage.

## Results

### Sequence elements of the programmed frameshift within *prfB* are hyper-conserved

To determine the prevalence of the *prfB* programmed frameshift motif, we analyzed the *prfB* sequences of 12,751 representative bacterial species genomes from the NCBI RefSeq database as annotated by NCBI (22). We identified the *prfB* sequence in each of these genomes and determined whether they contained a premature, in-frame stop codon within the *prfB* reading frame. Of the 12,751 genomes surveyed, 8160 (64%) contain a premature stop codon in *prfB*, indicating that these organisms require a programmed ribosomal frameshift to produce functional RF2.

We aligned the *prfB* sequences to generate a nucleotide sequence logo of the programmed frameshift motif (Fig. 2A). Within this motif, the purine-rich internal Shine-Dalgarno-like sequence is hyper-conserved, with the consensus sequence approaching “AGGGGG” (Fig. 2A). An even more highly conserved slippery sequence of “CTTT” occurs 3 nucleotides downstream of the internal Shine-Dalgarno-like sequence. The last T of the slippery sequence is the first T of the premature stop codon. The cytosine following the premature stop codon is also highly conserved, likely because “TGAC” causes poor termination efficiency and would permit more frequent frameshifting (14). This canonical “CTTTGA” sequence is present in nearly every *prfB* frameshift motif. Exceptions included 21 genomes with a long poly-thymine tract in the slippery sequence, in which tRNA_Phe_ would decode the codon before the stop codon instead of tRNA_Leu_ (i.e. “TTTTTGA”). These genomes belong to predominantly low-GC organisms (mean GC of 34.6%) in Aquificota and Gammaproteobacteria. In these genomes, the *prfB* sequence retains the internal SD-like sequence as well as the spacing between the SD-like sequence and the slippery sequence, and the identity of the premature stop codon is TGA.

**Figure 2.**
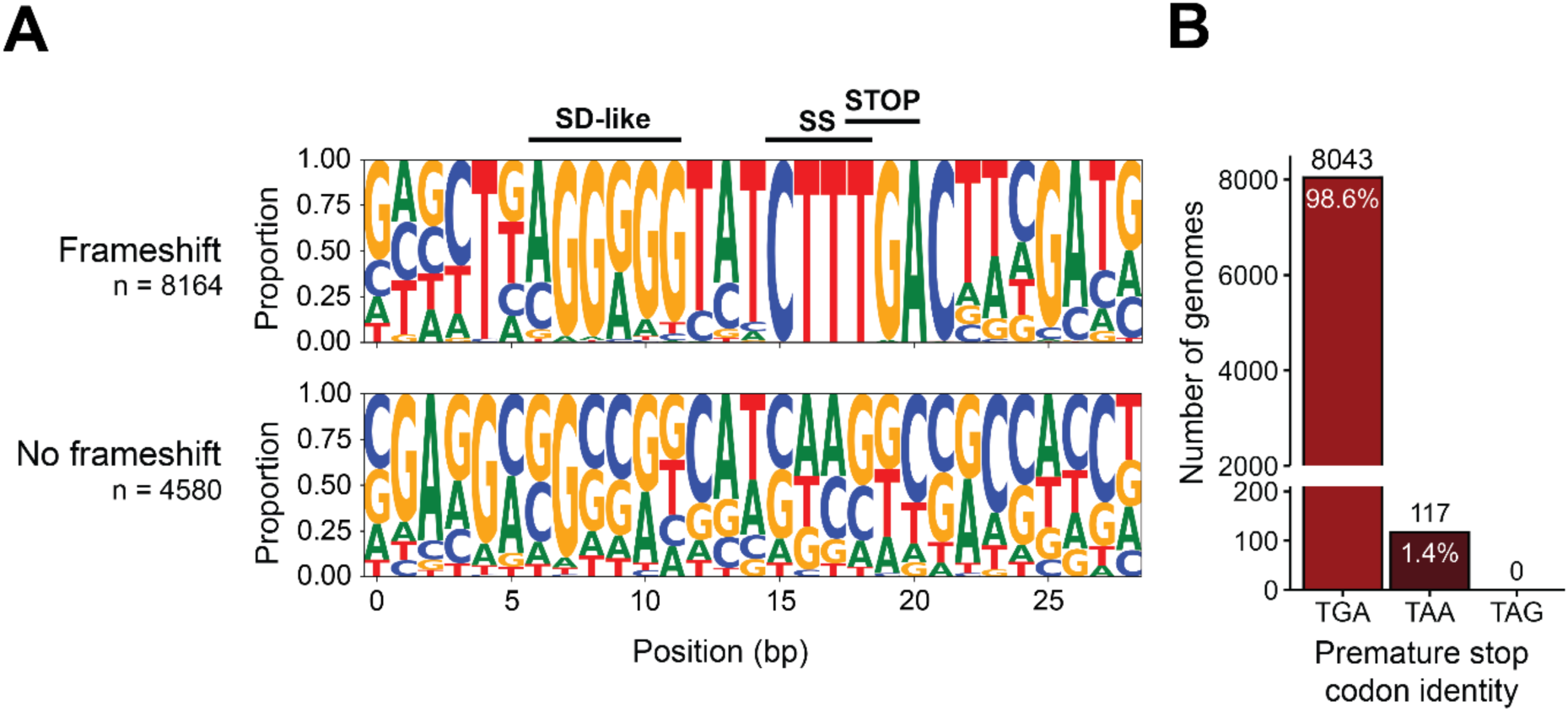
The premature stop codon in the *prfB* frameshifting motif is recognizable by RF2 in 100% of genomes surveyed. (A) Sequence logos of *prfB* frameshifting motif regions for genomes with and without a premature stop codon in *prfB*. The “No frameshift” logo captures the region in which the frameshifting motif is expected based on a full alignment of all *prfB* sequences. Labels above the black lines correspond to the following frameshifting motif components: SD-like, Shine-Dalgarno-like sequence; SS, slippery sequence; STOP, premature stop codon. (B) Identity of the premature stop codon within the *prfB* frameshifting motif.

We also aligned the *prfB* sequences of species that do not contain a premature stop codon to determine whether these sequences retained any elements of the programmed frameshift motif. In these *prfB* sequences the internal SD-like sequence and slippery sequence are not found in the analogous region of the sequence, suggesting that these elements are lost in organisms that encode a fully in-frame RF2 (Fig. 2A). This observation is expected since these elements would still promote ribosomal frameshifting, but without the presence of the autoregulatory TGA stop codon.

### The RF1-specific TAG stop codon is not detected as the premature stop codon in the *prfB* programmed frameshift motif

Bacteria terminate translation using one of three stop codons: TAA, TAG, or TGA. TAA is recognized by either RF1 or RF2, whereas TAG is RF1-specific and TGA is RF2-specific (23). To assess the conservation of the RF2-specific stop codon within the programmed frameshift motif, we determined the identity of the premature in-frame stop codon in the 8160 genomes that contain the motif. We found that 98.6% of genomes with the motif contain the RF2-specific TGA stop codon as the premature stop codon (“CTTTGA”) (Fig. 2B). 1.4% of genomes contain a TAA premature stop codon (“CTTTAA”). Since RF2 terminates translation at TAA as well as TGA, RF2 expression in these taxa would also be sensitive to the levels of RF2 in the cell. Genomes encoding TAA as the premature stop codon are observed randomly amongst phyla and retained among strains of a species (Fig. S1). These findings suggest that the TAA codon is poorly tolerated since it does not become fixed within particular clades and that the premature TGA is preferred for RF2 autoregulation (Fig. 2B). Most strikingly, none of the 8160 genomes encode an in-frame RF1-specific TAG stop codon. The ubiquity of TGA as the premature stop codon in the motif and the absence of TAG suggests that the universal purpose of the programmed frameshift motif is indeed RF2 autoregulation.

### The programmed frameshift *prfB* was likely present in the last common ancestor of bacteria

Bioinformatic analyses of the programmed frameshift motif in *prfB* sequences by Baranov and colleagues found that the motif is absent in *Aquifex aeolicus* and *Thermotoga maritime.* At the time, these taxa were thought to be the deepest branching taxa and most closely related to the ancestor of eubacteria. Therefore, absence of the programmed frameshift in the *prfB* of *A. aeolicus* and *T. maritime* suggested it may have been absent in the ancestral *prfB*. Recent efforts to root the bacterial tree position the root in the neighborhood of Fusobacteriota (24). This neighborhood also includes the Spirochaeotota, and a clade comprising Deinococcota, Synergistota, and Thermotogota (DST clade). Consistent with Baranov and colleagues, we did not detect the programmed frameshift in any of the 27 Thermotogota genomes we surveyed (Fig. 3A). However, we detected the programmed frameshift in >85% of *prfB* sequences from most other lineages in this neighborhood. The programmed frameshift is present in 87% of *prfB* sequences from Fusobacteriota (n = 31), 100% of Synergistota (n = 13), 99% of Deinococcota (n = 99), and 42% of Spirochaetota (n = 106) (Fig. 3A). The presence of the programmed frameshift motif in taxa closest to the root of the bacterial tree suggests that the ancestral *prfB* contained the programmed frameshift.

**Figure 3.**
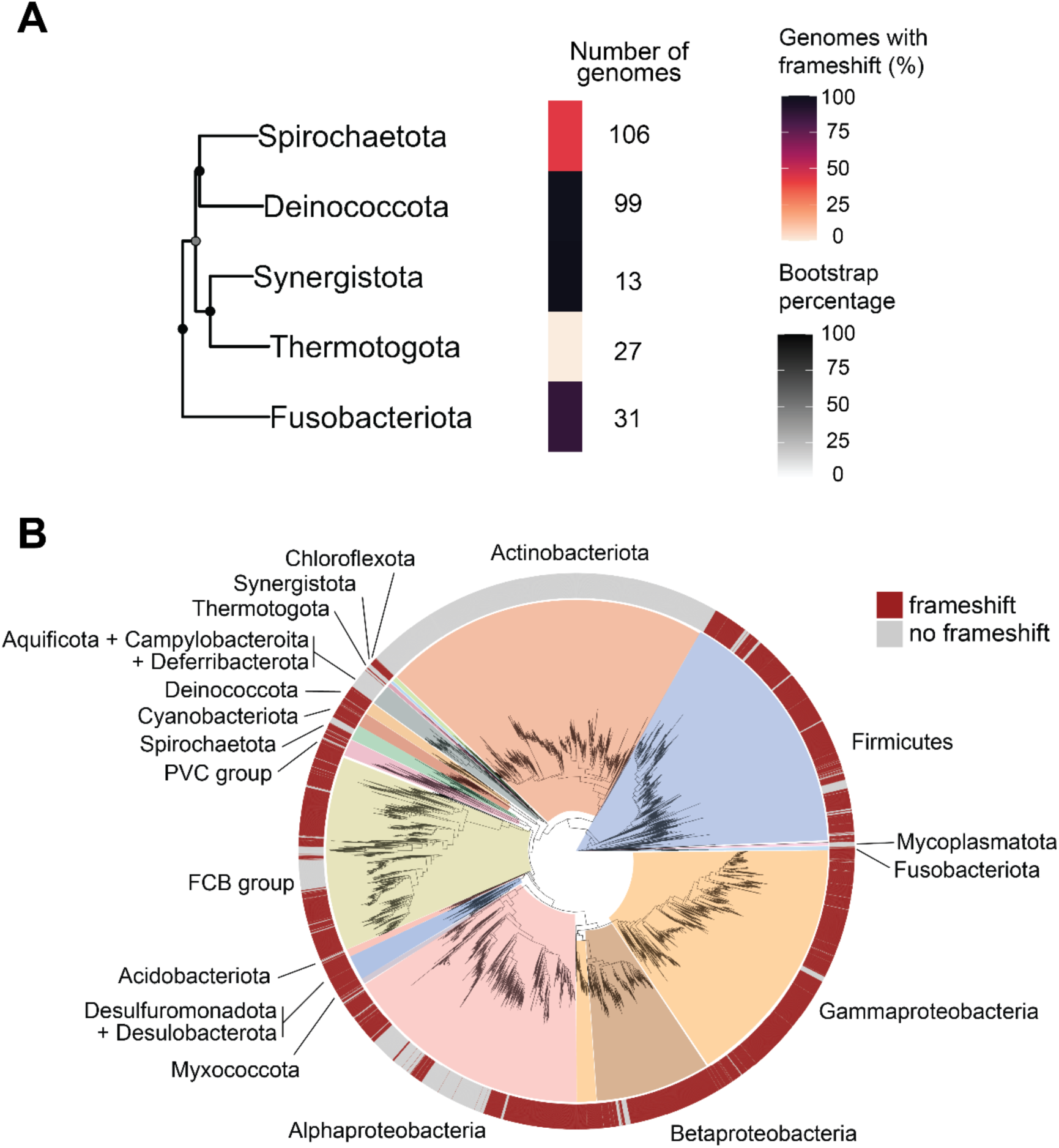
The *prfB* frameshifting motif is broadly distributed across bacterial phyla. (A) Percent of genomes containing the *prfB* frameshifting motif within phyla most closely related to the last bacterial common ancestor. A subtree of the large 16S tree was created using a random single representative genome for each phylum. The number of analyzed genomes per phylum is located to the right of each bar. Darkened circles indicate bootstrap values for each node. (B) A 16S maximum-likelihood phylogenetic tree showing distribution of genomes that encode the programmed frameshift within *prfB*. Phyla with more than 10 available and high-quality reference genomes are shown.

Next, we determined the distribution of the *prfB* programmed frameshift throughout the bacterial domain (Fig. 3B, Fig. S2). Only 3 phyla with >10 available genomes completely lacked the frameshift motif: Actinobacteriota, Mycoplasmatota, and Thermotogota. In 14 out of 19 phyla with >10 available genomes, >50% of genomes contain the premature stop and frameshifting motif (Fig. S2), demonstrating strong conservation of the programmed frameshift. For phyla where most genomes contain the motif, it is likely and parsimonious that the common ancestor of the phylum contained the motif, and that the motif was lost in recent lineages. Altogether, our results suggest that the programmed ribosomal frameshift in *prfB* is widespread throughout bacteria, including in deeply rooted branches of the bacterial domain.

### The *prfB* programmed frameshift motif is absent in Actinobacteriota and Mycoplasmatota

It was previously hypothesized that the lineages of Mycobacteria and Streptomyces lost the programmed frameshift in *prfB* based on its absence in two species – *Mycobacterium tuberculosis* and *Streptomyces coelicolor* (25). Expanding upon this, we surveyed the *prfB* sequences of the Actinobacteriota phylum that includes these lineages and did not detect the programmed frameshift in any of these genomes (n = 2658) (Fig. 3B), supporting a loss event in the common Actinobacterial ancestor.

Our analysis also indicates that the *prfB* programmed frameshift was likely absent in the common ancestor of the Mycoplasmatota phylum. Mycoplasmatota species are most closely related to Firmicutes (26), but have significantly reduced genome sizes, low GC content (27), and particularly low TGA stop codon usage (Fig. 4). Many species within Mycoplasmatota lack the *prfB* gene completely (28, 29) and utilize nearly zero TGA stop codons (12). Instead, TGA codons are decoded as tryptophan by suppressor tRNAs (27, 29–31). We found that all genomes lacking *prfB* encoded a suppressor tRNA for the TGA stop codon (Fig. 4). Only 23 out of 128 Mycoplasmatota genomes retained *prfB,* and none of these sequences contain the programmed frameshift motif (Fig. 4). Species that retain *prfB* are predominantly found within the Acholeplasmataceae family (Fig. 4), including important *Phytoplasma* plant pathogens (32).

**Figure 4.**
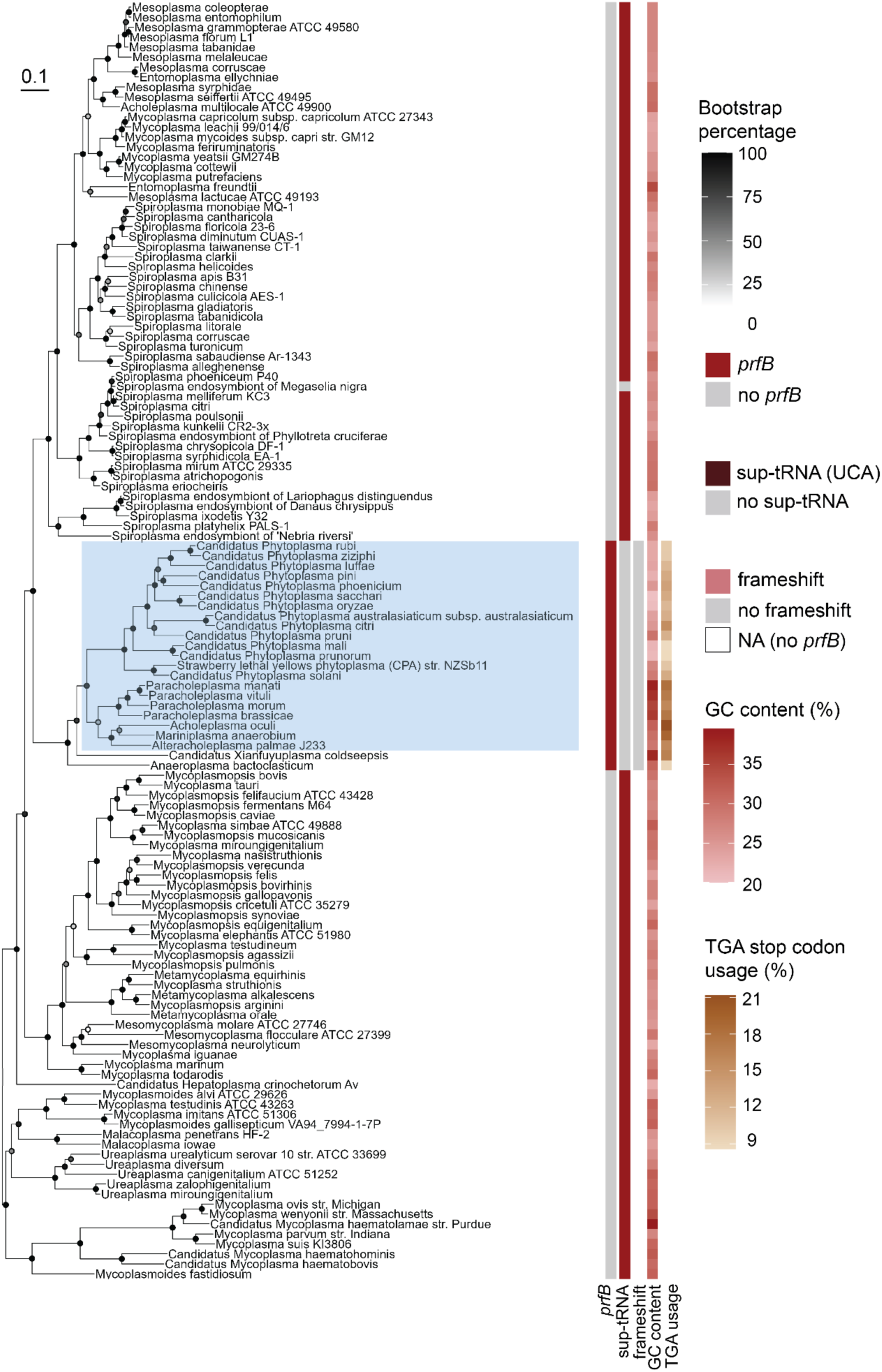
Most Mycoplasmatota species lack prfB and suppress TGA stop codons. Core SNP maximum-likelihood tree of 128 Mycoplasmatota species. From left to right, columns represent *prfB* gene presence, TGA stop codon suppressor tRNA presence, *prfB* frameshift motif presence, GC content, and TGA stop codon usage. The Acholeplasmataceae family is highlighted in blue. Darkened circles indicate bootstrap values for each node.

### Both free-living and obligate intracellular species retain the *prfB* programmed frameshift motif

Alphaproteobacteria are unique in that about 50% of genomes in this phylum encode the programmed ribosomal frameshift within *prfB* (Fig. S2). Since this phylum comprises both free-living and obligate intracellular bacteria, we questioned whether intracellular species would be less likely to autoregulate RF2 levels with the programmed ribosomal frameshift. We observed that Alphaproteobacterial genomes that do not use the frameshift motif in *prfB* fall predominantly in two orders: Rhodobacterales (n = 486 genomes) and Sphingomonadales (n = 338 genomes). Within Rhodobacterales, only two genomes appear to utilize the programmed frameshift motif. These were the only genomes of their respective genera, so additional genomes for the species could not be consulted to rule out sequence inaccuracy. Within Sphingomonadales, only 15 genomes contained the frameshifting motif, and these genomes were in the family Sphingosinicellaceae. Members of both Rhodobacterales and Sphingomonadales tend to be free-living and are found in soils, water, and sediments (33). Other free-living orders of the Alphaproteobacteria such as Caulobacterales and Rhodospirillales predominantly encode the programmed frameshift motif in *prfB*. Meanwhile, the obligate intracellular Alphaproteobacterial genera including *Rickettsia* and *Wolbachia* maintain the *prfB* gene and its frameshifting motif. Therefore, we conclude that the programmed frameshift motif is conserved in both free-living and in obligate intracellular species within the Alphaproteobacteria.

### Genomes that lack the programmed frameshifting motif in *prfB* have higher GC content and more TGA stop codon usage

We next explored genome characteristics of organisms that do not utilize the programmed frameshift in *prfB*. We found that genomes that lack the programmed frameshift motif have significantly higher GC content than genomes with the motif (62% average GC, p < 2.2e-16) (Fig. 5A). Our finding remains significant even with the removal of the well-represented high-GC Actinobacterial genomes (p < 1.1e-07) (Fig. 5B). GC content positively correlates with RF2-specific TGA codon usage (12) (Fig. S3). Therefore, we hypothesized that organisms lacking RF2 autoregulation would also encode more RF2-specific stop codons. To investigate this, we compared terminal stop codon usage between genomes with and without the programmed frameshift for a random subset of 1000 genomes. Genomes that lost the *prfB* frameshift motif have significantly higher RF2-specific TGA terminal stop codon usage than genomes that retained the motif (p < 2.2e-16) (Fig. 5A). Again, our findings are significant even when Actinobacterial genomes are excluded (p < 2.2 e-06) (Fig. 5B). Therefore, a higher demand for RF2 due to increased RF2-specific TGA stop codon usage may help explain the loss of the RF2 autoregulation in some species.

**Figure 5.**
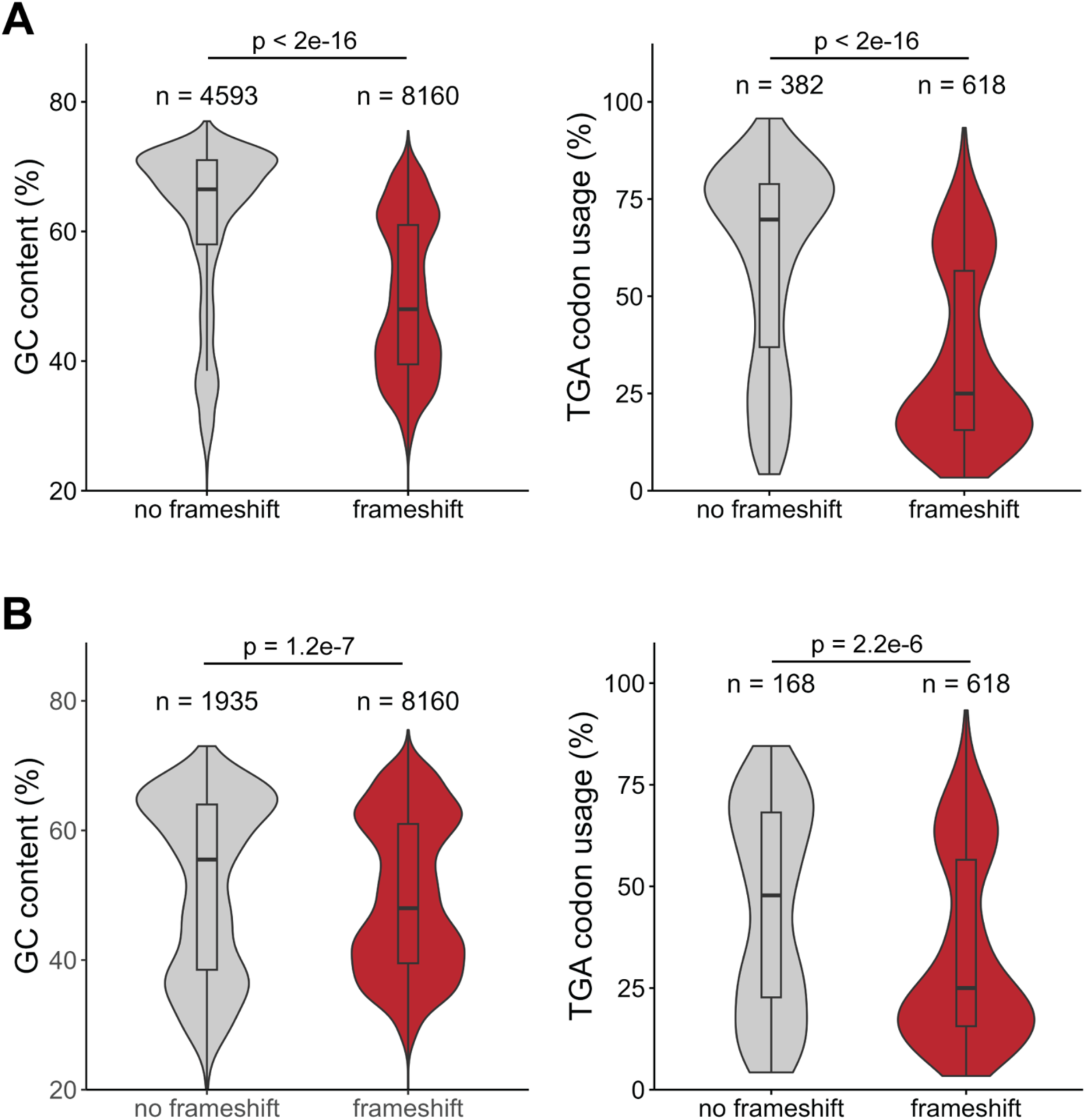
Genomes that encode the programmed frameshift motif within *prfB* have significantly higher GC content and TGA stop codon usage. (A) GC content and TGA stop codon usage of a subset of 1000 random genomes separated by genomes with and without the *prfB* frameshift motif. (B) Distributions of GC content and TGA stop codon usage of the random subset without Actinobacterial genomes are also shown. P-values indicate the results of a Welch two-sample *t*-test.

### Ribosomal frameshifting is inefficient at the *prfB* programmed frameshift motif in the Actinobacterium *Mycobacterium smegmatis*

No surveyed Actinobacterial genomes (n = 2658) contain the *prfB* frameshift. Therefore, we hypothesized that Actinobacteriota may exhibit poor frameshift efficiency at the motif. To assay frameshifting efficiency, we compared the frameshifting efficiencies of an organism that natively contains the frameshift motif, *B. subtilis*, and an Actinobacterium that lacks the motif, *Mycobacterium smegmatis*. We designed analogous constructs for *B. subtilis* and *M. smegmatis* that contain two fused protein sequences (encoding mCherry and GFP) separated by the *prfB* frameshift motif from *B. subtilis* (denoted as ‘FS motif’). We included ∼60 bp upstream and downstream of the frameshifting motif to ensure that the context surrounding the frameshift motif is also maintained. In *M. smegmatis*, the fluorescent proteins and the 60 bp regions upstream and downstream of the *prfB* motif were codon optimized for *M. smegmatis* to avoid ribosomes stalling at rare codons. As a control for production of the truncated protein we used a construct identical to the *prfB* FS motif construct including the premature TGA stop codon but lacking the frameshifting motif elements (denoted as ‘no FS motif’). Construct schematics are depicted in Fig. 6A.

**Figure 6.**
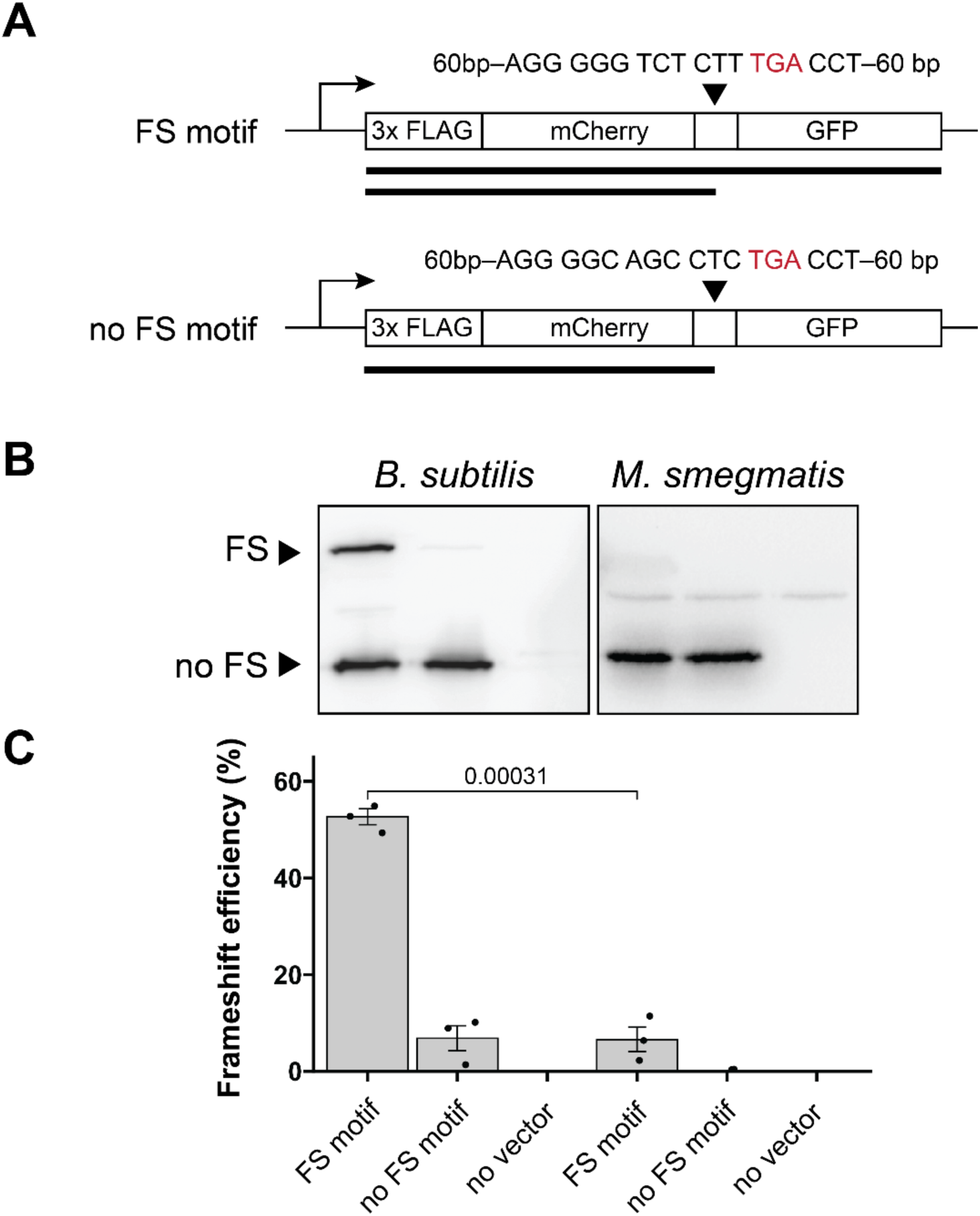
*Mycobacterium smegmatis* cannot efficiently frameshift at the canonical *prfB* frameshifting motif. (A) Schematics for analogous prfB frameshifting reporters in *B. subtilis* and *M. smegmatis*. (B) Western blots showing both frameshifted and termination products resulting from expression of the pictured constructs. Blots are representative of three biological replicates. (C) Quantification of western blots. Frameshifting efficiency = frameshifted protein produced/(frameshifted + non-frameshifted protein produced). P-values indicate the results of a Welch two-sample *t*-test.

Upon expressing the previously described constructs *in vivo*, we observed that ribosomal frameshifting occurs at the *prfB* motif with an efficiency of 52.7% ± 2.8% in *B. subtilis* (Fig. 6B and 6C). In contrast, *M. smegmatis* exhibits ribosomal frameshifting at the motif with an efficiency of 6.6% ± 4.4% (Fig. 6B and 6C). As expected, neither organism exhibits strong ribosomal frameshifting at the *prfB* variant that lacks the frameshifting motif elements (Fig. 6B). The abnormally low frameshift efficiency in *M. smegmatis* at the *prfB* motif supports the idea that members of the Actinobacteriota have lost this form of RF2 autoregulation because they are unable to efficiently utilize the programmed frameshifting motif.

## Discussion

The autoregulation of *prfB* expression by RF2 levels is a textbook example of programmed ribosomal frameshifting, and the molecular mechanism underlying the programmed frameshift has been precisely characterized in *E. coli* (11, 13). Using >12,000 bacterial genomes and referencing a recently updated bacterial phylogenetic tree (24), we present evidence that this regulatory frameshift motif was likely present in the ancestral *prfB*. We find that loss of the programmed frameshift highly correlates with RF2 specific stop codon usage. This finding is consistent with the expectation that the requirement for higher RF2 levels will select for inactivation of an autoregulatory mechanism that reduces RF2 expression.

Our analysis of the programmed frameshifting elements in *prfB* is consistent with previous reports analyzing fewer than 100 genomes (19, 21, 25) and is in perfect agreement with the highly characterized mechanistic features of the ribosomal frameshift. In particular, the SD-like sequence, the spacing between the SD-like sequence and the premature stop codon, the slippery sequence preceding the stop codon, and the C nucleotide immediately following the stop codon are almost universally conserved. Most striking, the identity of the premature stop codon is TGA in >99% of surveyed genomes and is TAA in all remaining genomes surveyed. There is evidence that TAA may be primarily recognized by RF2 in most bacterial taxa (34–38), with the exception of *E. coli* K-12 strains which encode a less efficient RF2 (36). Therefore, even *prfB* genes containing a premature TAA stop codon would likely be subject to regulation by RF2 levels. The precise conservation of each of the frameshifting elements in taxa throughout the bacterial domain suggests that nearly all bacteria that autoregulate RF2 levels do so by the same molecular mechanism.

Previous work postulated that the ancestral *prfB* lacked the programmed ribosomal frameshift. This hypothesis was informed by an understanding of the bacterial tree as being rooted closest to Thermotogota and the absence of the programmed frameshift in *prfB* sequences from this phylum. We similarly did not detect the ribosomal frameshift in *prfB* sequences in Thermotogota (Fig. 3A). However, the overwhelming majority of *prfB* sequences of all phyla in proximity to Thermotogota do indeed utilize the programmed frameshift (Fig. 3A). Because current models place the root of the bacterial tree in the neighborhood of Fusobacteria, and the majority of taxa in this neighborhood, except Thermotogota, use the programmed frameshift in *prfB*, it is likely and parsimonious that the ancestral *prfB* was regulated by a programmed ribosomal frameshift.

Having determined that the ancestral *prfB* likely encoded the programmed frameshift, we next examined the genome characteristics of taxa that lost the frameshift motif. We found that bacteria that have lost the programmed frameshift in *prfB* had significantly higher RF2-specific stop codon usage (p < 2.2e^-16^) (Fig. 5A). Even when we excluded Actinobacteriota, which make up a large proportion of genomes that do not encode the motif, this finding was still highly significant (p = 2.2e^-06^) (Fig. 5B). Bacterial release factor concentrations correlate with cognate stop codon usage (12, 39). The direction of causality is unknown, but a recent *in silico* study proposed that release factor concentrations adapted to stop codon usage (40). High GC content strongly correlates with high RF2-specific TGA stop codon usage but not with TAG stop codon usage (Fig S3) (12). Therefore, a likely explanation for RF2 autoregulation loss is that high GC content and high RF2-specific stop codon usage increased the demand for RF2, and RF2 autoregulation was subsequently lost to satisfy this demand.

Across the bacterial tree of life, the most notable loss event occurred in the Actinobacterial ancestor. None of the 2658 Actinobacterial genomes in our survey encoded the programmed frameshift in *prfB*. Therefore, we hypothesized that Actinobacteriota may exhibit poor ribosomal frameshifting efficiency at this motif. In support of this hypothesis, when we provided *M. smegmatis* with a reporter encoding the *prfB* frameshift motif, the frameshift efficiency was ∼6%. We cannot conclude from this result whether *M. smegmatis* ribosomes are less able to frameshift at this frameshift motif due to structural differences, or whether RF2 levels and activity in *M. smegmatis* are high enough that termination at the UGA stop codon is favored over frameshifting. In either case, these results suggest that encoding the frameshift motif in *prfB* would be detrimental in *M. smegmatis* because it may result in decreased RF2 levels. Interestingly, the only other organism reported to have similarly low frameshifting efficiency at the *prfB* motif is *Flavobacterium johnsoniae,* which rarely uses Shine-Dalgarno sequences during translation initiation (41, 42). Surprisingly, *F. johnsoniae* retains the *prfB* frameshift motif despite its low frameshifting efficiency. The low efficiency may be tolerated because the *F. johnsoniae* genome has an unusually low proportion of RF2-specific stop codons (7% of all stop codons) (41), and therefore a low demand for RF2.

Although RF2 specific stop codon usage is highly correlated with loss of the motif, there are notable exceptions. In particular, Mycoplasmatota have low genomic GC content and low TGA stop codon usage, but Mycoplasmatota genomes that encode *prfB* all lack the frameshifting motif (Fig. 4). The ancestral Mycoplasmatota underwent extreme genome reduction, and therefore, many species in the phylum have lost *prfB,* instead decoding UGA as tryptophan (29, 43). It is possible that in lineages that retain *prfB*, RF2 may be actively decaying and therefore lowly expressed, even without autoregulation. Therefore, RF2 autoregulation is not required in these lineages.

Our work supports a model in which the programmed ribosomal frameshift motif in *prfB* was present in the last common ancestor of bacteria and autoregulates RF2 expression in most bacterial species. In many species that have lost the motif, it is likely that high TGA stop codon usage increased demand for RF2 and so RF2 autoregulation is no longer required. While our work offers a comprehensive survey of the evolution and purpose of RF2 autoregulation, future studies are needed to determine whether there are release factor or structural differences in the ribosome that also affect frameshifting efficiency in diverse bacteria and may have selected for loss of the *prfB* programmed frameshifting motif.

### Data Availability Statement

Data for all genomes surveyed can be found in Table S1 and S2. Scripts for data acquisition and analyses are available on GitHub at https://github.com/cassprince/prfB_evolution.

## Materials and Methods

### Strains and media

All strains were derived from *B. subtilis* 168 *trpC2* and *M. smegmatis* MC^2^ 155. *B. subtilis* was grown shaking in Lysogeny Broth media at 37°C, and *M. smegmatis* was grown shaking in 7H9 Middlebrook media at 37°C as indicated. Antibiotics were used at final concentrations of 5 µg/mL chloramphenicol and 20 µg/mL kanamycin. Plasmids used in this study are described in Table 1 and novel plasmid sequences are available on the project GitHub at https://github.com/cassprince/prfB_evolution/blob/main/data/plasmid_sequences.fasta. All experiments were performed in biological triplicate.

**Table 1.**
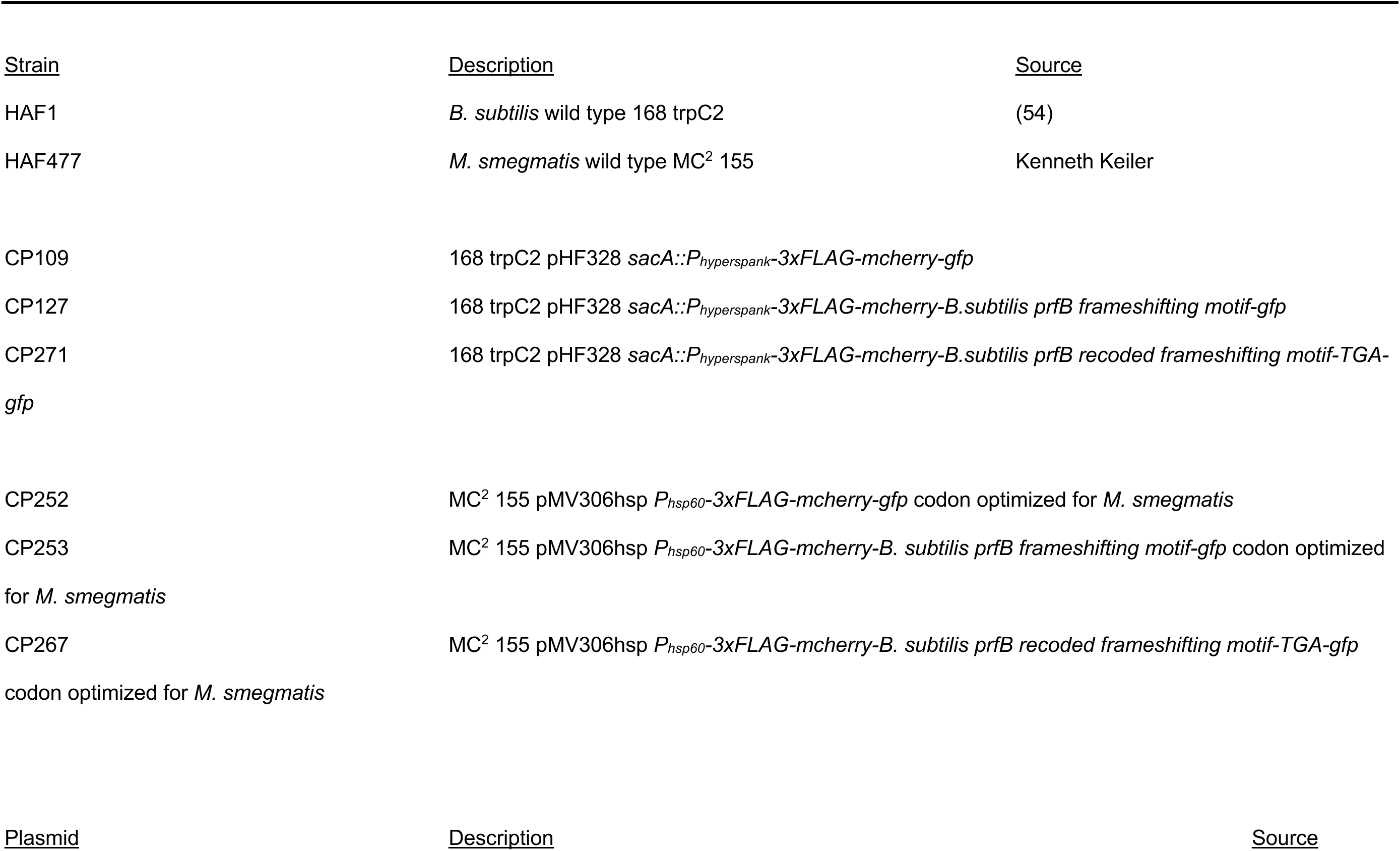
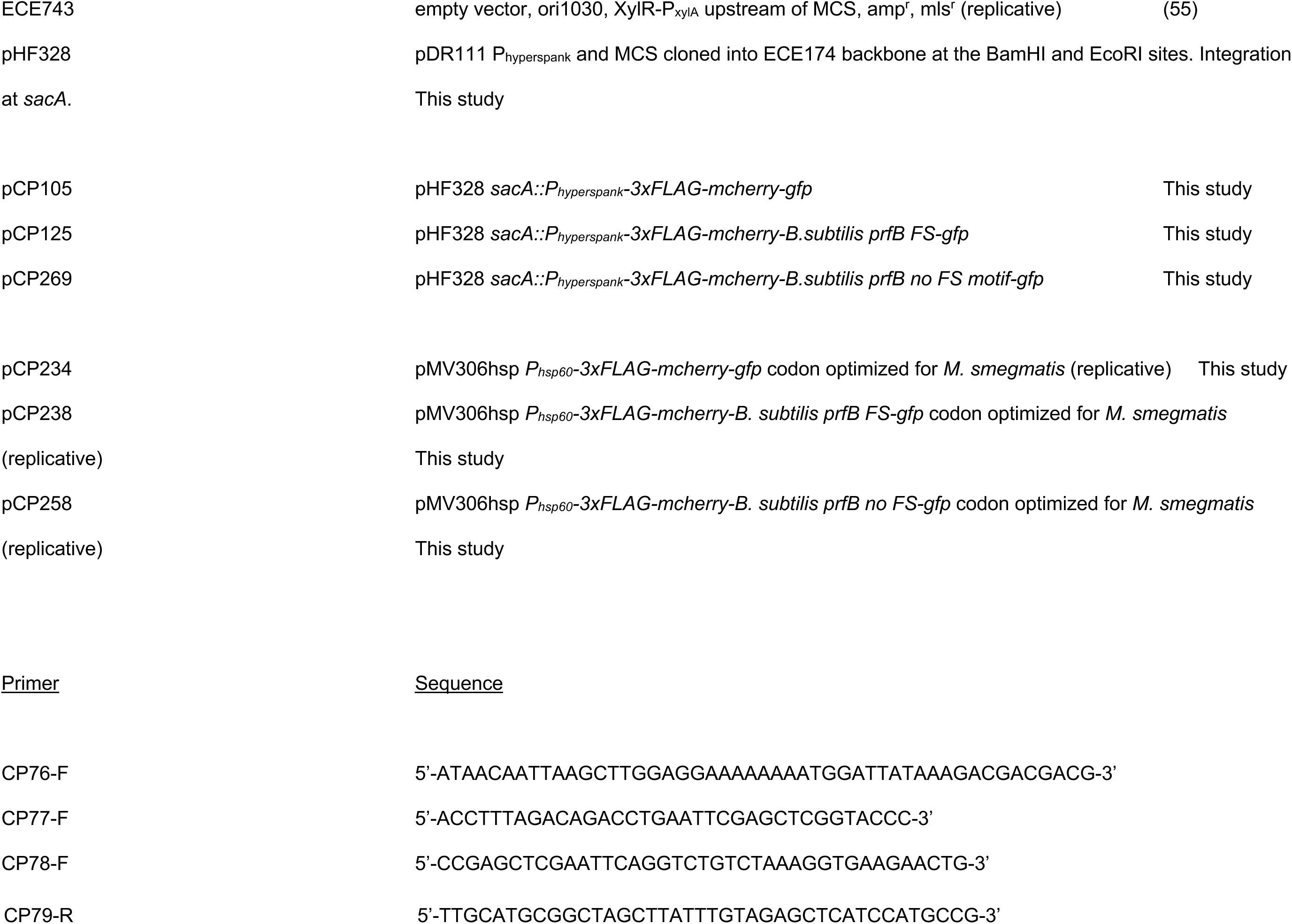
Strains, Plasmids, and Primers.

### Frameshift reporter lysates and western blots

*M. smegmatis* overnight cultures were normalized to an OD600 of 0.2. Cells were then harvested and pelleted after 12 hours. *B. subtilis* overnight cultures were normalized to an OD600 of 0.05. Reporter expression was induced with 1mM IPTG when cultures reached an OD600 of 1. Cells were then harvested and pelleted after 30 minutes of induction. *B. subtilis* and *M. smegmatis* cell pellets were treated with lysis buffer (10 mM Tris pH 8, 50 mM EDTA, 1 mg/mL lysozyme) for 10 minutes at 37°C. *M. smegmatis* cells were further lysed using bead beating for five cycles of 20 seconds at 4350 rpm with 3 minutes on ice between cycles. All lysates were mixed with SDS loading dye, boiled at 90°C for 5 minutes, and cooled on ice. For *M. smegmatis*, the protein levels of the full-length reporter were approximately 15x greater than those of the experimental *prfB* frameshifting reporters based on band intensity. Therefore, the lysates for the full-length reporter strain were diluted 15x to normalize the protein levels between reporters.

Proteins were run on a 12% SDS-PAGE gel for 70 minutes at 150 V. To measure *prfB* overexpression levels and reporter frameshifting levels, proteins were transferred from SDS-PAGE gels to a PVDF membrane (BioRad) for 100 minutes at 300 mAmps. The membrane was blocked in 3% bovine serum albumin (BSA) overnight at 4°C. Anti-FLAG antibody conjugated to horseradish peroxidase (Sigma SAB4200119) was added to the BSA for 1.5 hours at room temperature. The membrane was washed with PBS-T three times for 5 minutes at room temperature and developed with ECL substrate and enhancer (Biorad 170-5060). Band intensities were quantified using ImageJ v1.53k (44). P-values for differences in band intensity were calculated with the R stats v4.2.2 package using a Welch two-sampled *t*-test.

### *prfB* sequence acquisition and analyses

*prfB* nucleotide sequences were downloaded from all representative prokaryotic genomes in the NCBI RefSeq database as annotated by the NCBI Prokaryotic Genome Annotation Pipeline (22). Only genomes with CheckM (45) contamination scores below 10% were used, resulting in 12,751 genomes. The NCBI accession numbers, species names, and taxids for all genomes used can be found in Table S1. *prfB* sequences were aligned using MAFFT v7.453 (46). Python scripts were used to identify premature stop codons by searching for any stop codon that was in-frame but not found in the final three nucleotides of the sequence. The surrounding region was then extracted, and the identity of the stop codon was recorded. If no premature stop codon was found, the expected region of the frameshifting motif (based on multiple sequence alignment) was extracted. The scripts utilized the biopython v1.78 (47) package for sequence manipulation. The extracted regions were converted to sequence logos using the logomaker v0.8 package (48).

### Phylogenetic analyses

16S rRNA sequences were identified and acquired using BLAST v2.13.0. Sequences were aligned using MAFFT v7.453 (46). The alignments were applied to FastTree v2.1.11 (49) to infer a maximum likelihood tree. Trees were visualized using the ggtree v3.6.2 package (50). FastTree produces unrooted phylogenies, so trees were midpoint rooted using the phangorn v2.11.1 package (51). The simplified tree in Figure 3A was produced by randomly selecting a representative genome for each phylum and subsetting the large 16S tree in Figure 3B. The identities of the randomly selected genomes can be found in Table S3. Taxonomic classification was assigned to genomes using the NCBI Taxonomy database (52) and taxonkit v0.14.1 (53). GC content for each genome was downloaded from NCBI. To determine terminal stop codon usage, all coding sequences were downloaded as annotated by NCBI PGAP for a random subset of 1000 genomes. The list of genomes in the subset can be found in Table S2. A novel Python script recorded the last three nucleotides of each coding sequence per genome, utilizing the biopython v1.78 package for sequence manipulation. P-values for differences in GC content and terminal stop codon usage between “frameshift” and “no frameshift” genomes were calculated with the R stats v4.2.2 package using a Welch two-sample t-test.

## Acknowledgements

HAF and CRP were supported by NIH R35GM147049. CRP was supported by a Graduate Research Fellowship from the National Science Foundation.

## Supplemental Figures

**Figure S1.**
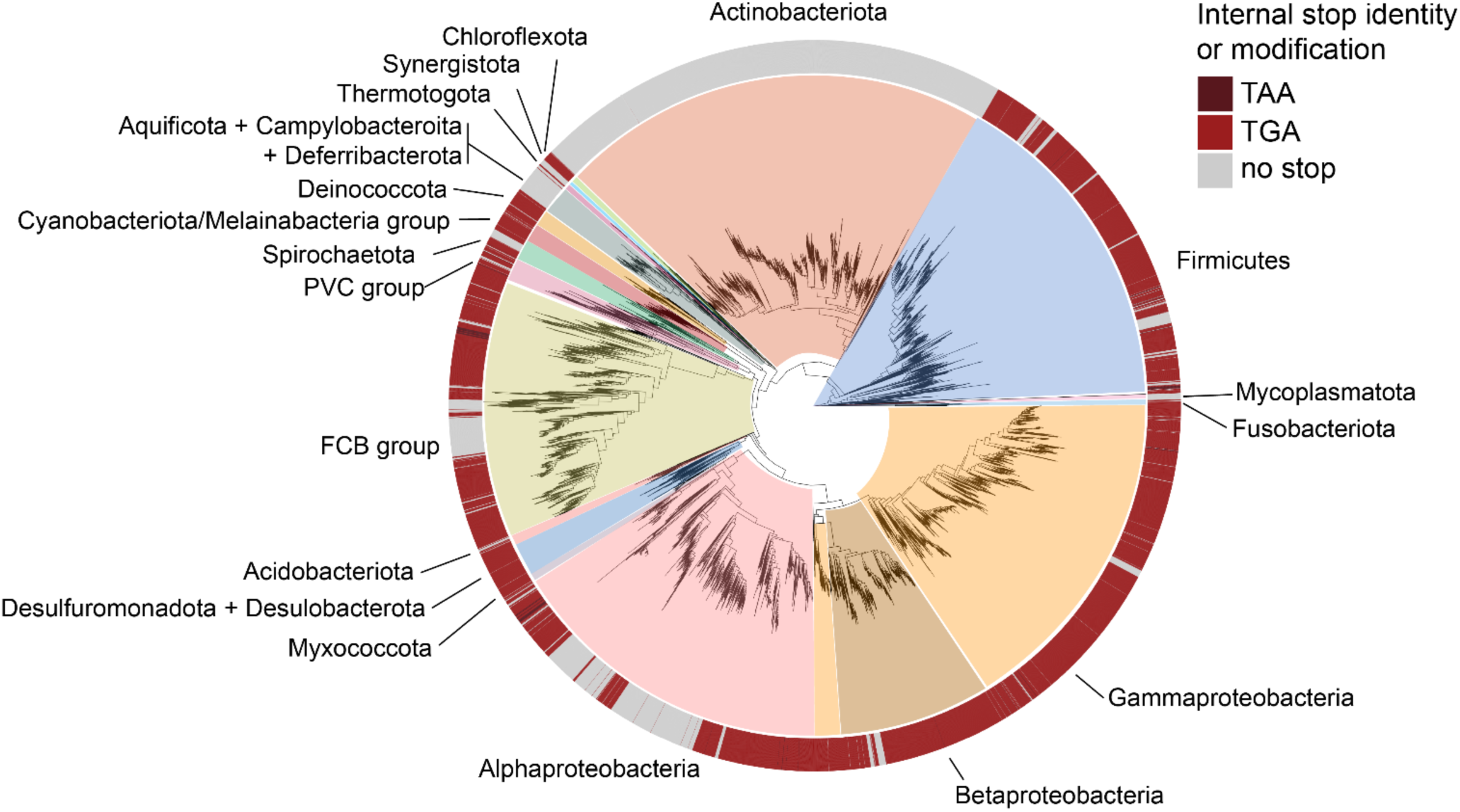
TAA is poorly represented and randomly distributed as the premature stop codon in the frameshifting motif. 16S maximum-likelihood phylogenetic tree from Figure 3B showing the distribution of premature stop codons in *prfB*. Phyla with more than 10 available and high-quality reference genomes are shown.

**Figure S2.**
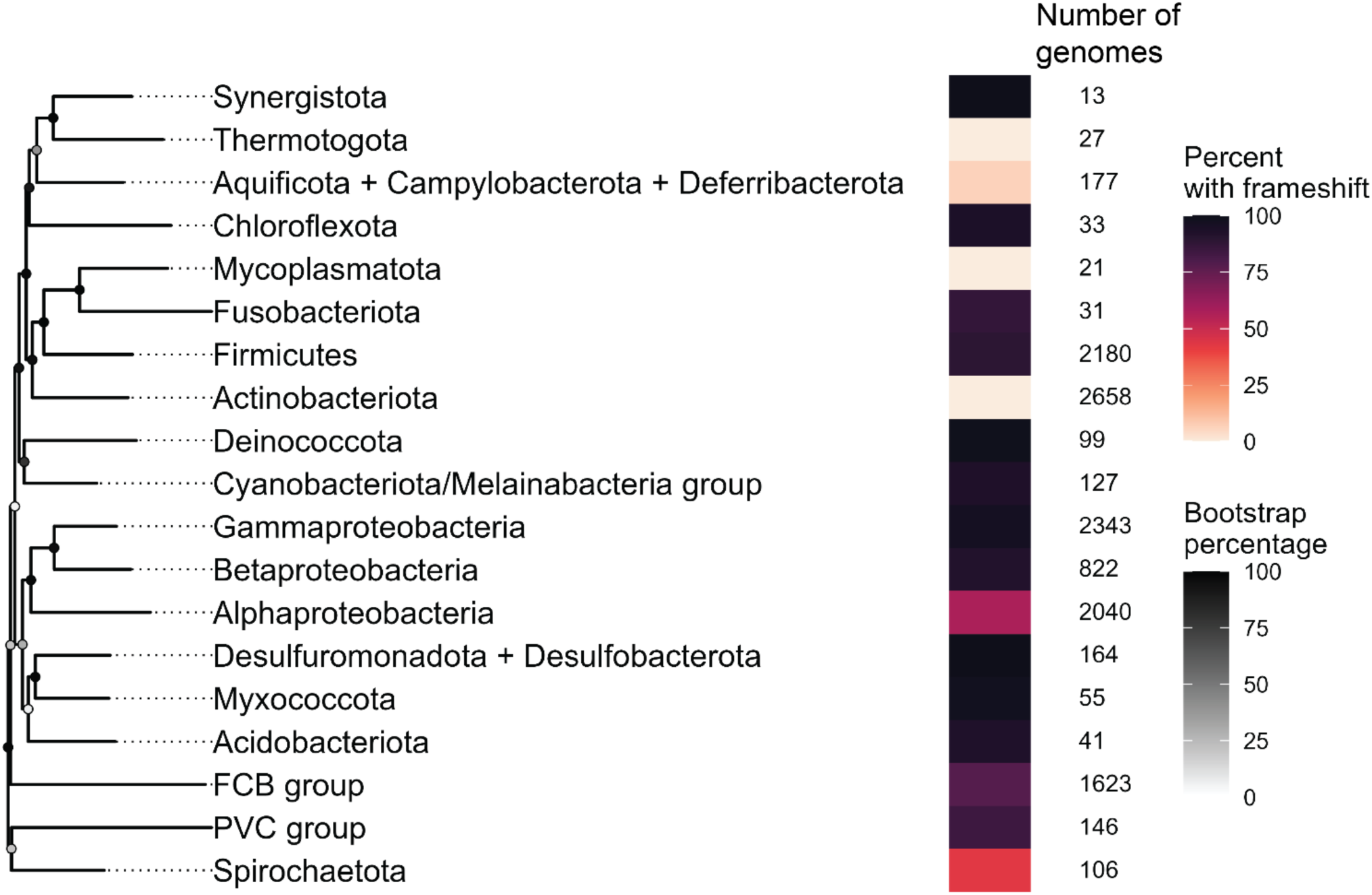
Percent of genomes containing the *prfB* frameshifting motif across phyla. A subtree of the large 16S tree was created using a random single representative genome for each phylum with more than 10 genomes. The number of analyzed genomes per phylum is located to the right of each bar. Darkened circles indicate bootstrap values for each node.

**Figure S3.**
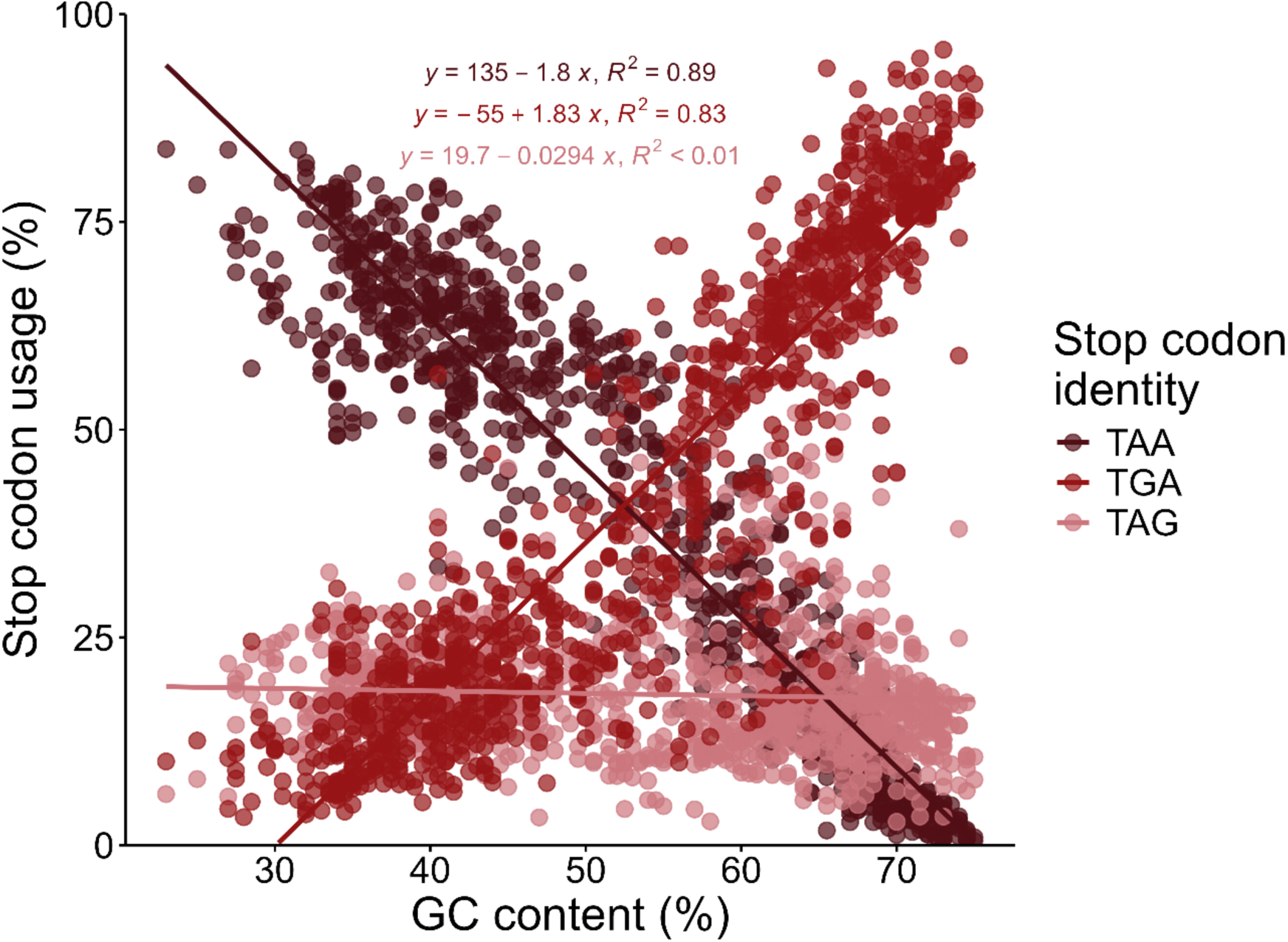
TGA stop codon usage increases with GC content. Terminal stop codon usage in all coding sequences for a subset of 1000 random reference genomes plotted verses GC content. Each genome has one point for each of the three stop codons.

## Supplemental table legends

**Table S1.** Metadata of the 12,571 genomes used in this study.

**Table S2.** Total stop codon usage for the random subset of 1,000 genomes in Figure 5.

**Table S3.** Accession numbers for random representatives of phyla in Figure 3A and S2.

## Notes

https://github.com/cassprince/prfB_evolution

### Competing Interest Statement

The authors have declared no competing interest.

### Summary of Updates

This version has been revised to focus on the computational aspects of the work.

## References

1. Tse H, Cai JJ, Tsoi H-W, Lam EP, Yuen K-Y. 2010. Natural selection retains overrepresented out-of-frame stop codons against frameshift peptides in prokaryotes. BMC Genomics 11:491.

2. Atkins JF, Gesteland RF, Reid BR, Anderson CW. 1979. Normal tRNAs promote ribosomal frameshifting. Cell 18:1119–1131.

3. Kastelein RA, Remaut E, Fiers W, van Duin J. 1982. Lysis gene expression of RNA phage MS2 depends on a frameshift during translation of the overlapping coat protein gene. Nature 295:35–41.

4. Larsen B, Gesteland RF, Atkins JF. 1997. Structural probing and mutagenic analysis of the stem-loop required for Escherichia coli dnaX ribosomal frameshifting: programmed efficiency of 50%11Edited By D. E. Draper. 1. Journal of Molecular Biology 271:47–60.

5. Meydan S, Klepacki D, Karthikeyan S, Margus T, Thomas P, Jones JE, Khan Y, Briggs J, Dinman JD, Vázquez-Laslop N, Mankin AS. 2017. Programmed Ribosomal Frameshifting Generates a Copper Transporter and a Copper Chaperone from the Same Gene. 2. Molecular Cell 65:207–219.

6. Farabaugh PJ. 1996. Programmed translational frameshifting. Annu Rev Genet 30:507–528.

7. Dinman JD. 2012. Control of gene expression by translational recoding. Adv Protein Chem Struct Biol 86:129–149.

8. Dinman JD. 2012. Mechanisms and implications of programmed translational frameshifting. Wiley Interdiscip Rev RNA 3:661–673.

9. Advani VM, Dinman JD. 2016. Reprogramming the genetic code: The emerging role of ribosomal frameshifting in regulating cellular gene expression. Bioessays 38:21–26.

10. Craigen WJ, Cook RG, Tate WP, Caskey CT. 1985. Bacterial peptide chain release factors: conserved primary structure and possible frameshift regulation of release factor 2. Proc Natl Acad Sci U S A 82:3616–3620.

11. Craigen WJ, Caskey CT. 1986. Expression of peptide chain release factor 2 requires high-efficiency frameshift. 6076. Nature 322:273–275.

12. Korkmaz G, Holm M, Wiens T, Sanyal S. 2014. Comprehensive Analysis of Stop Codon Usage in Bacteria and Its Correlation with Release Factor Abundance *. 44. Journal of Biological Chemistry 289:30334–30342.

13. Weiss RB, Dunn DM, Atkins JF, Gesteland RF. 1987. Slippery runs, shifty stops, backward steps, and forward hops: -2, -1, +1, +2, +5, and +6 ribosomal frameshifting. Cold Spring Harb Symp Quant Biol 52:687–693.

14. Poole ES, Brown CM, Tate WP. 1995. The identity of the base following the stop codon determines the efficiency of in vivo translational termination in Escherichia coli. 1. EMBO J 14:151–158.

15. Major LL, Poole ES, Dalphin ME, Mannering SA, Tate WP. 1996. Is the in-frame termination signal of the Escherichia coli release factor-2 frameshift site weakened by a particularly poor context? Nucleic Acids Res 24:2673–2678.

16. Poole ES, Major LL, Mannering SA, Tate WP. 1998. Translational termination in Escherichia coli: three bases following the stop codon crosslink to release factor 2 and affect the decoding efficiency of UGA-containing signals. Nucleic Acids Res 26:954–960.

17. Márquez V, Wilson DN, Tate WP, Triana-Alonso F, Nierhaus KH. 2004. Maintaining the ribosomal reading frame: the influence of the E site during translational regulation of release factor 2. Cell 118:45–55.

18. Weiss RB, Dunn DM, Dahlberg AE, Atkins JF, Gesteland RF. 1988. Reading frame switch caused by base-pair formation between the 3ʹ end of 16S rRNA and the mRNA during elongation of protein synthesis in Escherichia coli. The EMBO Journal 7:1503–1507.

19. Baranov PV, Gesteland RF, Atkins JF. 2002. Recoding: translational bifurcations in gene expression. Gene 286:187–201.

20. Baranov PV, Gesteland RF, Atkins JF. 2002. Release factor 2 frameshifting sites in different bacteria. 4. EMBO Rep 3:373–377.

21. Bekaert M, Atkins JF, Baranov PV. 2006. ARFA: a program for annotating bacterial release factor genes, including prediction of programmed ribosomal frameshifting. 20. Bioinformatics 22:2463–2465.

22. Li W, O’Neill KR, Haft DH, DiCuccio M, Chetvernin V, Badretdin A, Coulouris G, Chitsaz F, Derbyshire MK, Durkin AS, Gonzales NR, Gwadz M, Lanczycki CJ, Song JS, Thanki N, Wang J, Yamashita RA, Yang M, Zheng C, Marchler-Bauer A, Thibaud-Nissen F. 2020. RefSeq: expanding the Prokaryotic Genome Annotation Pipeline reach with protein family model curation. D1. Nucleic Acids Res 49:D1020–D1028.

23. Scolnick E, Tompkins R, Caskey T, Nirenberg M. 1968. Release factors differing in specificity for terminator codons. Proc Natl Acad Sci U S A 61:768–774.

24. Coleman GA, Davín AA, Mahendrarajah TA, Szánthó LL, Spang A, Hugenholtz P, Szöllősi GJ, Williams TA. 2021. A rooted phylogeny resolves early bacterial evolution. 6542. Science 372:eabe0511.

25. Persson BC, Atkins JF. 1998. Does disparate occurrence of autoregulatory programmed frameshifting in decoding the release factor 2 gene reflect an ancient origin with loss in independent lineages? 13. J Bacteriol 180:3462–3466.

26. Hug LA, Baker BJ, Anantharaman K, Brown CT, Probst AJ, Castelle CJ, Butterfield CN, Hernsdorf AW, Amano Y, Ise K, Suzuki Y, Dudek N, Relman DA, Finstad KM, Amundson R, Thomas BC, Banfield JF. 2016. A new view of the tree of life. Nat Microbiol 1:1–6.

27. Sirand-Pugnet P, Citti C, Barré A, Blanchard A. 2007. Evolution of mollicutes: down a bumpy road with twists and turns. Research in Microbiology 158:754–766.

28. Fraser CM, Gocayne JD, White O, Adams MD, Clayton RA, Fleischmann RD, Bult CJ, Kerlavage AR, Sutton G, Kelley JM, Fritchman RD, Weidman JF, Small KV, Sandusky M, Fuhrmann J, Nguyen D, Utterback TR, Saudek DM, Phillips CA, Merrick JM, Tomb JF, Dougherty BA, Bott KF, Hu PC, Lucier TS, Peterson SN, Smith HO, Hutchison CA, Venter JC. 1995. The minimal gene complement of Mycoplasma genitalium. Science 270:397–403.

29. Grosjean H, Breton M, Sirand-Pugnet P, Tardy F, Thiaucourt F, Citti C, Barré A, Yoshizawa S, Fourmy D, Crécy-Lagard V de, Blanchard A. 2014. Predicting the Minimal Translation Apparatus: Lessons from the Reductive Evolution of Mollicutes. PLOS Genetics 10:e1004363.

30. Inamine JM, Ho KC, Loechel S, Hu PC. 1990. Evidence that UGA is read as a tryptophan codon rather than as a stop codon by Mycoplasma pneumoniae, Mycoplasma genitalium, and Mycoplasma gallisepticum. J Bacteriol 172:504–506.

31. Citti C, Maréchal-Drouard L, Saillard C, Weil JH, Bové JM. 1992. Spiroplasma citri UGG and UGA tryptophan codons: sequence of the two tryptophanyl-tRNAs and organization of the corresponding genes. J Bacteriol 174:6471–6478.

32. Doi Y, Teranaka M, Yora K, Asuyama H. 1967. Mycoplasma- or PLT Group-like Microorganisms Found in the Phloem Elements of Plants Infected with Mulberry Dwarf, Potato Witches’ Broom, Aster Yellows, or Paulownia Witches’ Broom. Japanese Journal of Phytopathology 33:259–266.

33. Hördt A, López MG, Meier-Kolthoff JP, Schleuning M, Weinhold L-M, Tindall BJ, Gronow S, Kyrpides NC, Woyke T, Göker M. 2020. Analysis of 1,000+ Type-Strain Genomes Substantially Improves Taxonomic Classification of Alphaproteobacteria. Front Microbiol 11:468.

34. Mora L, Heurgué-Hamard V, de Zamaroczy M, Kervestin S, Buckingham RH. 2007. Methylation of bacterial release factors RF1 and RF2 is required for normal translation termination in vivo. J Biol Chem 282:35638–35645.

35. Elliott T, Wang X. 1991. Salmonella typhimurium prfA mutants defective in release factor 1. J Bacteriol 173:4144–4154.

36. Uno M, Ito K, Nakamura Y. 1996. Functional specificity of amino acid at position 246 in the tRNA mimicry domain of bacterial release factor 2. Biochimie 78:935–943.

37. Baggett NE, Zhang Y, Gross CA. 2017. Global analysis of translation termination in E. coli. PLoS Genet 13:e1006676.

38. Dinçbas-Renqvist V, Engström Å, Mora L, Heurgué-Hamard V, Buckingham R, Ehrenberg M. 2000. A post-translational modification in the GGQ motif of RF2 from Escherichia coli stimulates termination of translation. EMBO J 19:6900–6907.

39. Wei Y, Wang J, Xia X. 2016. Coevolution between Stop Codon Usage and Release Factors in Bacterial Species. Molecular Biology and Evolution 33:2357–2367.

40. Ho AT, Hurst LD. 2022. Variation in Release Factor Abundance Is Not Needed to Explain Trends in Bacterial Stop Codon Usage. Molecular Biology and Evolution 39:msab326.

41. Naeem FM, Gemler BT, McNutt ZA, Bundschuh R, Fredrick K. 2024. Analysis of programmed frameshifting during translation of prfB in Flavobacterium johnsoniae. 2. RNA 30:136–148.

42. McNutt ZA, Gandhi MD, Shatoff EA, Roy B, Devaraj A, Bundschuh R, Fredrick K. 2021. Comparative Analysis of anti-Shine-Dalgarno Function in Flavobacterium johnsoniae and Escherichia coli. Front Mol Biosci 8:787388.

43. Yamao F, Iwagami S, Azumi Y, Muto A, Osawa S, Fujita N, Ishihama A. 1988. Evolutionary dynamics of tryptophan tRNAs in Mycoplasma capricolum. Mol Gen Genet 212:364–369.

44. Schneider CA, Rasband WS, Eliceiri KW. 2012. NIH Image to ImageJ: 25 years of image analysis. Nat Methods 9:671–675.

45. Parks DH, Imelfort M, Skennerton CT, Hugenholtz P, Tyson GW. 2015. CheckM: assessing the quality of microbial genomes recovered from isolates, single cells, and metagenomes. Genome Res 25:1043–1055.

46. Katoh K, Standley DM. 2013. MAFFT Multiple Sequence Alignment Software Version 7: Improvements in Performance and Usability. 4. Molecular Biology and Evolution 30:772–780.

47. Cock PJA, Antao T, Chang JT, Chapman BA, Cox CJ, Dalke A, Friedberg I, Hamelryck T, Kauff F, Wilczynski B, de Hoon MJL. 2009. Biopython: freely available Python tools for computational molecular biology and bioinformatics. 11. Bioinformatics 25:1422–1423.

48. Tareen A, Kinney JB. 2020. Logomaker: beautiful sequence logos in Python. 7. Bioinformatics 36:2272–2274.

49. Price MN, Dehal PS, Arkin AP. 2010. FastTree 2 – Approximately Maximum-Likelihood Trees for Large Alignments. 3. PLOS ONE 5:e9490.

50. Yu G, Smith DK, Zhu H, Guan Y, Lam TT-Y. 2017. ggtree: an r package for visualization and annotation of phylogenetic trees with their covariates and other associated data. 1. Methods in Ecology and Evolution 8:28–36.

51. Schliep KP. 2011. phangorn: phylogenetic analysis in R. 4. Bioinformatics 27:592–593.

52. Schoch CL, Ciufo S, Domrachev M, Hotton CL, Kannan S, Khovanskaya R, Leipe D, Mcveigh R, O’Neill K, Robbertse B, Sharma S, Soussov V, Sullivan JP, Sun L, Turner S, Karsch-Mizrachi I. 2020. NCBI Taxonomy: a comprehensive update on curation, resources and tools. Database (Oxford) 2020:baaa062.

53. Shen W, Ren H. 2021. TaxonKit: A practical and efficient NCBI taxonomy toolkit. Journal of Genetics and Genomics 48:844–850.

54. Gaidenko TA, Kim T-J, Price CW. 2002. The PrpC Serine-Threonine Phosphatase and PrkC Kinase Have Opposing Physiological Roles in Stationary-Phase Bacillus subtilis Cells. J Bacteriol 184:6109–6114.

55. Popp PF, Dotzler M, Radeck J, Bartels J, Mascher T. 2017. The Bacillus BioBrick Box 2.0: expanding the genetic toolbox for the standardized work with Bacillus subtilis. Sci Rep 7:15058.

